# Fast P(RMNE): Fast Forensic DNA Probability of Random Man Not Excluded Calculation

**DOI:** 10.1101/173708

**Authors:** Darrell O. Ricke, Steven Schwartz

**Affiliations:** Bioengineering Systems & Technologies Massachusetts Institute of Technology Lincoln Laboratory Lexington, MA USA

**Keywords:** DNA forensic, identification, mixture analysis

## Abstract

High throughput sequencing (HTS) of DNA forensic samples is expanding from the sizing of short tandem repeats (STRs) to massively parallel sequencing (MPS). HTS panels are expanding from the FBI 20 core Combined DNA Index System (CODIS) loci to include SNPs. The calculation of random man not excluded, P(RMNE), is used in DNA mixture analysis to estimate the probability that a person is present in a DNA mixture. This calculation encounters calculation artifacts with expansion to larger panel sizes. Increasing the floating-point precision of the calculations allows for increased panel sizes but with a corresponding increase in computation time. The Taylor series higher precision libraries used fail on some input data sets leading to algorithm unreliability. Herein, a new formula is introduced for calculating P(RMNE) that scales to larger SNP panel sizes while being computationally efficient (patent pending)[1].

## I. INTRODUCTION

High throughput sequencing (HTS) of DNA single nucleotide polymorphism (SNP) panels have significant advantages for analysis of DNA mixtures and trace DNA profiles compared to sizing STRs. Analysis of mixtures by sized STRs is limited to mixtures of two individuals within DNA ratios of 1:1 to 1:10. In contrast, SNP-based methods offer the potential to analyze complex mixtures of 15 contributors or more[2]. The current method of calculating the significance of a match between a SNP DNA mixture and a reference profile is the random man not excluded P(RMNE) calculation[2] for forensic applications. However, performance and precision issues are being observed with current implementations of the P(RMNE) calculations[2]. To address the calculation artifacts and performance issues, a novel P(RMNE) calculation method is presented.

## II. METHODS

### A. Taylor series P(RMNE) implementation

Most SNPs have just two alleles. The most common SNP allele is named the major allele. The other SNP allele(s) are named the minor allele(s). In a mixture profile, the minor allele ratio is calculated as the ratio of minor allele reads divided by the total number of reads. Methods for calculating P(RMNE) have been presented that focus on the mixture SNP loci with no called minor alleles in a mixture profile (e.g., SNPs with minor allele ratios <= 0.001 threshold)[2, 3]. The P(RMNE) method described by Isaacson et al.[2] was implemented in Sherlock’s Toolkit[4]. This formulation enabled P(RMNE) calculations with a small number of dropped alleles for reference profiles compared to mixture profiles. For larger DNA panels, an issue with precision was observed with the Sherlock’s Toolkit implementation, see Figure 1. This method was re-implemented in Java with higher precision libraries in an effort to eliminate the calculation artifacts observed (Figure 1). The Discrete Fourier Transform-Characteristic Function (DFT-CF) method was implemented with Taylor series approximation of trigonometric functions, named Taylor-32 for 32-bit floating point and Taylor-64 for 64-bit floating point calculations.

**Figure 1.**
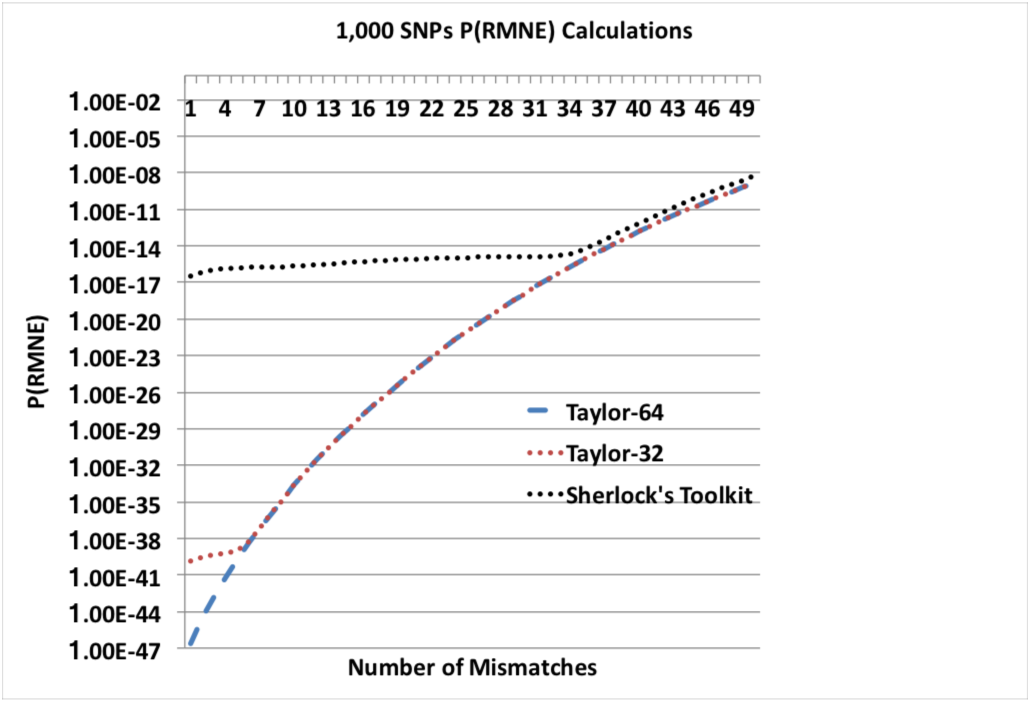
P(RMNE) Results for 1,000 SNP Panel.

### B. Mathar’s BigDecimalMath P(RMNE) calculation

The Taylor series library functions were replaced with functions from Mathar’s BigDecimalMath class[5] to address issues detected with the Taylor-32 and Taylor-64 methods using both 64-bit and 152-bit precision.

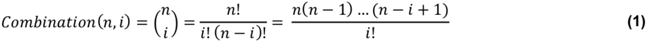

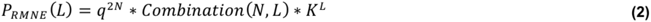

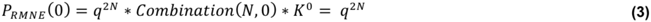

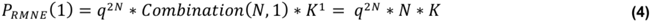

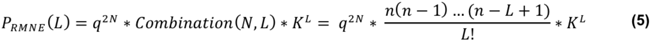

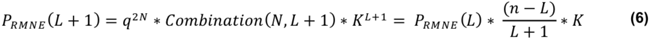

**Figure 2.**
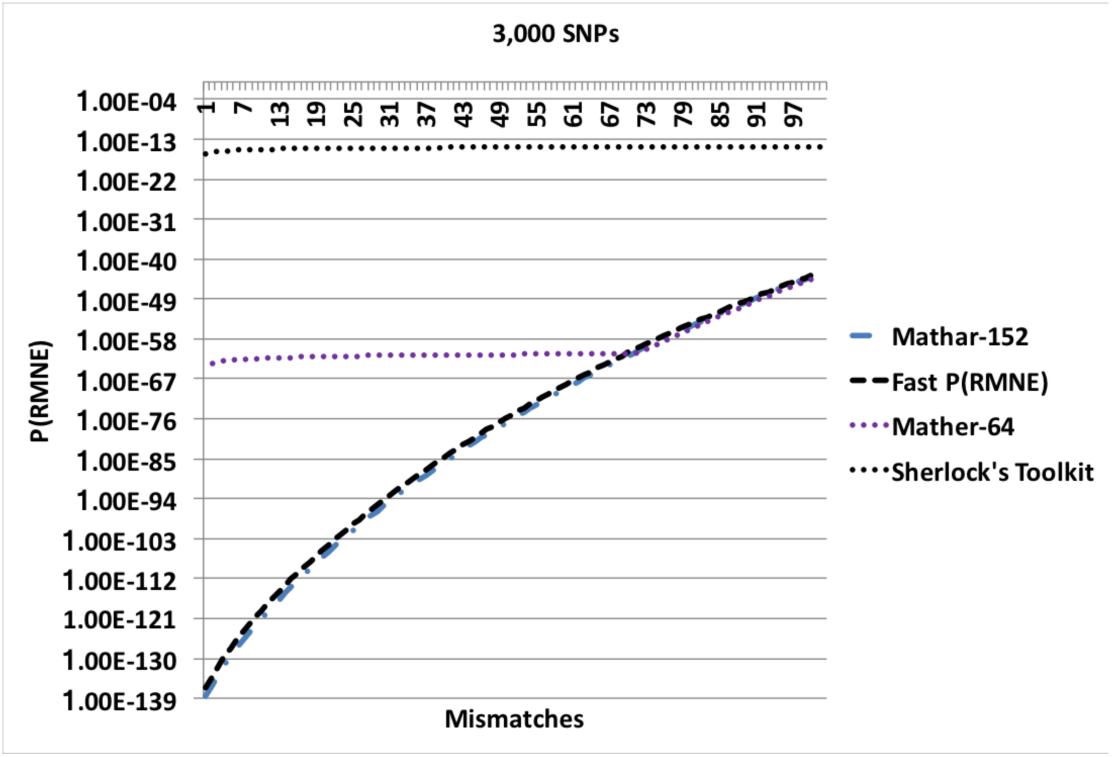
P(RMNE) Results for 3,000 SNP Panel.

### C. Fast P(RMNE)

An alternative to the DFT-CF P(RMNE) method was implemented. A mixture will have N loci with no called minor alleles. Let p be the average minor allele ratio at these mixture loci. Let q be defined as 1 – p such that p + q = 1. SNP panels can be optimized for DNA mixture analysis[2, 3]; the average of the SNP minor allele ratios used for a P(RMNE) calculation can be used to approximate large numbers of individual SNPs with similar minor allele ratios. For an individual with two alleles at a SNP loci the probability for these alleles can be represented as (p+q)^2^ = p^2^ + 2pq + q^2^ = 1. A perfect reference match to a mixture has major:major (MM) alleles at every locus with no called minor alleles in the mixture profile. Mismatches are defined as reference loci with major:minor (mM) or minor:minor (mm) at these mixture loci with no called minor alleles (MM). The number of mismatches is defined as L between a reference and a mixture. Let K be (1 – q^2^)/q^2^ represent the ratio of transition from MM to non-MM (i.e., mM or mm). Let Combination represent the standard statistics combination operation for representing possible SNP loci that mismatch between a reference and a mixture (1). P_RMNE_(L) can be estimated by the term for no mismatches, q^2N^, times the possible combinations of L mismatches, Combination(N, L), times the transition term K^L^ (2)[2]. Equation (3) illustrates the calculation for no mismatches (L=0), and (4) for one mismatch (L=1). Consecutive terms can be calculated efficiently for multiple L values as illustrated by (5) and (6). This optimization has the additional benefit of multiplying a large value, (N-L)/(L+1), with a small value, K, where calculating N!/L!(N-L)! by itself can stress the precision capability of an implementation for large values for N and L. Equation (7) represents the P(RMNE) calculation for 0 to L mismatches.

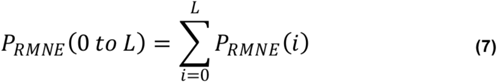

### D. Benchmark Systems

Timing for the Sherlock’s Toolkit (Python), Taylor, and Mathar algorithms (Java) were run on an Intel Xeon E5-2609 v2 2.5 GHz dual CPU system with 32 GB RAM. Fast P(RMNE) (Ruby) was run on a MacBook Pro laptop with 2.8 GHz Intel i7, 16 GB 1600 MHz DDR3 RAM, 750 GB SSD hard drive.

## III. RESULTS

The calculated P(RMNE) values for Sherlock’s Toolkit and Taylor-32 both have calculation artifacts/precision issues compared to the Taylor-64 method for a panel of 1,000 SNPs in Figure 1. The Sherlock’s Toolkit P(RMNE) values start to deviate from actual P(RMNE) values with 36 or less mismatches while the Taylor-32 deviates at 5 or less mismatches. When the panel size is increased to 3,000 SNPs, the Taylor methods are unable to calculate P(RMNE) values. For higher precision, the Mathar BigDecimalMath library was used with 64-bit and 152-bit precision. Calculation artifacts are seen for the Mathar 64-bit method for the 3,000 SNP panel (Figure 2) and the Mathar 152-bit method for the 4,000 SNP panel (Figure 3). The root mean square error (RMSE) between Fast P(RMNE) and Mathar-152 was 2.2e-41. This calculation excluded the Mathar 152-bit calculation artifacts between 0 and 19 mismatches. Algorithm timing results are shown in Figure 4. For the 1,000 SNP panel, the Taylor 64-bit algorithm runs in 142 s and the Taylor 152-bit in 1,017 s. The Taylor methods did not complete for the larger panel sizes.

**Figure 3.**
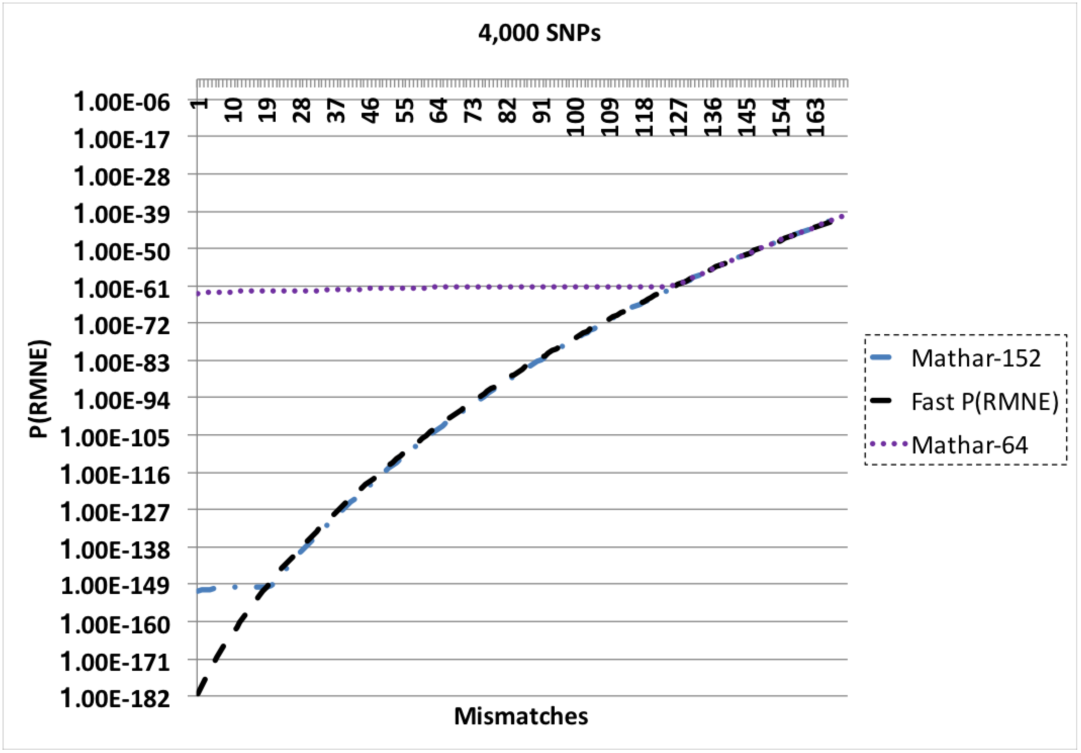
P(RMNE) Results for 4,000 SNP Panel.

**Figure 4.**
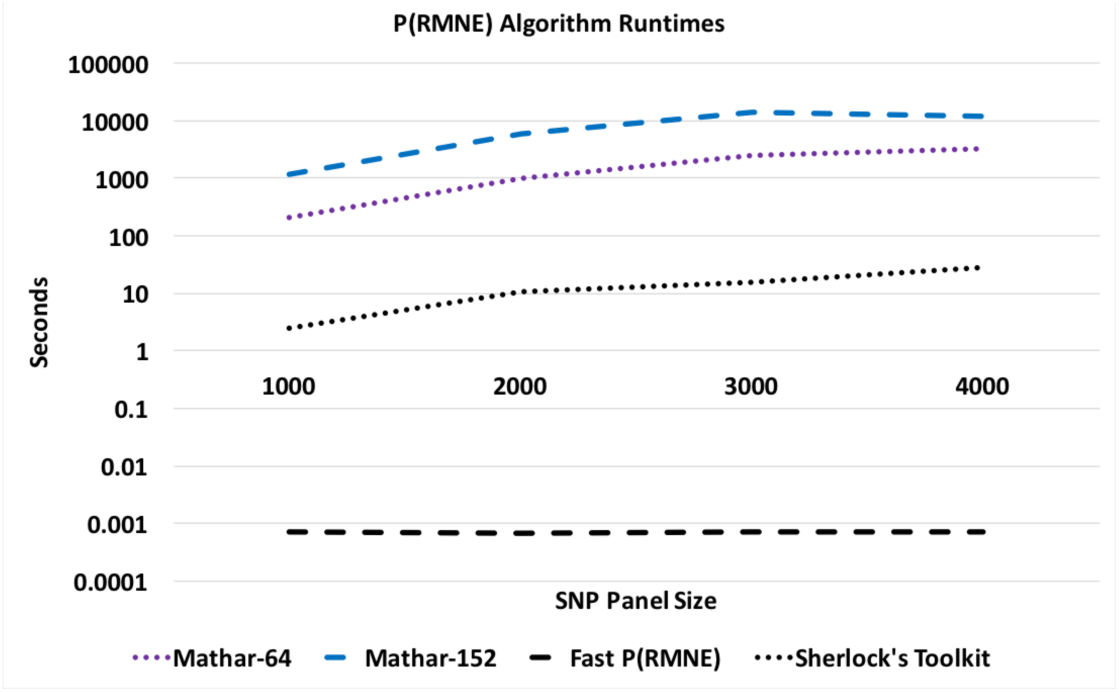
P(RMNE) Algorithm Runtimes.

## IV. DISCUSSION

A calculation artifact was observed for some datasets with the P(RMNE) method implemented in Sherlock’s Toolkit, see Figure 1. Shifting to higher precision libraries, improved the results for smaller SNP panels, but calculation artifacts appear for larger SNP panels, see Figures 2 and 3. Also, the Taylor methods crash with larger panels or return no results. The Mathar BigDecimalMath libraries work better than the Taylor method library, but calculation artifacts are again observed for the 4,000 SNP panels for both Mathar-64 and Mathar-152 methods. The runtimes for these higher precision methods as increased beyond what was desirable for rapid forensic sample analysis. The Fast P(RMNE) method addresses both the calculation artifact issue (Figure 3) and the runtime issue (Figure 4). Equation (6) enables the rapid calculation of P(RMNE) for a series of possible mismatches in a fraction of a second on any modern CPU processor.

## V. DISTRIBUTION STATEMENT

A. Approved for public release: distribution unlimited.

## ACKNOWLEDGMENT

We would like to thank Nancy DeLosa for assisting with the Taylor and Mathar methods implementations.

